# Common facultative endosymbionts do not influence sensitivity of cereal aphids to pyrethroids

**DOI:** 10.1101/2022.05.24.493255

**Authors:** D J Leybourne, P Melloh, E A Martin

## Abstract

1. Cereal aphids, including the bird cherry-oat aphid, *Rhopalosiphum padi*, and the grain aphid, *Sitobion avenae*, can transmit viruses that significantly reduce crop yields. To mitigate against yield losses, insecticides are routinely used to manage aphid populations.
2. Aphids can form relationships with endosymbionts that confer fitness benefits or consequences to the aphid. Recent artificial inoculation experiments indicate that endosymbionts could increase aphid susceptibility to insecticides, but this has not been explored using aphid populations naturally infected with endosymbionts.
3. Here, we sampled aphids from an important cereal production region in Lower Saxony, Germany. We characterised the endosymbiont profile of these aphid populations and conducted pyrethroid dose-response assays to test the hypothesis that facultative endosymbionts increase aphid susceptibility to insecticides.
4. We find that the level of insecticide susceptibility is highly variable in *S. avenae* and we identify populations that are sensitive and tolerant to pyrethroids, including populations collected from the same field. For *R. padi*, we find evidence for decreased sensitivity to pyrethroids, representing the first report of reduced sensitivity to pyrethroids in *R. padi* sampled from Central Europe.
5. We detected high endosymbiont infection frequencies in the aphid populations. 84% of aphids carry one facultative endosymbiont and 9% of aphids carry two facultative endosymbionts. We detected associations with *Regiella insecticola, Fukatsia symbiotica*, and *Hamiltonella defensa*. However, we do not identify a link between endosymbiont infection and insecticide susceptibility, indicating that other factors may govern the development of insecticide resistance and the need for alternative management strategies.

## Introduction

Cereal aphids, including the bird cherry-oat aphid, *Rhopalosiphum padi*, and the grain aphid, *Sitobion avenae*, are important herbivorous insects. Cereal aphids are classed as agricultural pest species on many grasses and cereals, including wheat and barley (Van Emden & Harrington, 2017). Cereal aphids are widely distributed across Central Europe and can cause significant damage to cereal crops. Aphid damage can be caused through direct feeding (Dedryver *et al*., 2010) and via the transmission of plant viruses, including barley yellow dwarf virus (BYDV) (Perry *et al*., 2000). High levels of BYDV infection in cereal crops can result in yield losses of *c*. 80% (Nancarrow *et al*., 2021).

Insecticides remain the method that is most commonly used to manage aphid populations, with pyrethroids widely used for the management of cereal aphids on spring and winter cereal crops across Europe (Dewar & Foster, 2017). The high reliance on pyrethroid insecticides increases the evolutionary pressure on aphid populations, increasing the risk that insecticide resistant aphid populations will emerge (Dewar & Foster, 2017). Insecticide resistant populations can have devastating consequences on effective aphid management and increase potential aphid-derived yield loss (Dewar & Foster, 2017), making these an urgent priority for the development of alternative management strategies. Resistance to insecticides evolves over time, and monitoring surveys of herbivorous insect populations can detect the emergence of insecticide resistance by identifying populations that are less sensitive to insecticides (Umina *et al*., 2020; Walsh *et al*., 2020a). Resistance against pyrethroids has been described in both *S. avenae* and *R. padi* populations (Foster *et al*., 2014; Wang *et al*., 2020). Pyrethroid resistance is associated with mutations in voltage-gated ion channels (Foster *et al*., 2014; Wang *et al*., 2020) and this resistance mechanism is referred to as *knock down resistance* (*kdr*). Two mutations conferring *kdr* resistance have been described, namely *kdr* (Foster *et al*., 2014) and *super-kdr* (Wang *et al*., 2020).

Pyrethroid resistance has been described in *S. avenae* populations, with heterozygous *knockdown resistance* (*kdr*-SR) resistant populations detected in China, Ireland, and the UK (Foster *et al*., 2014; Walsh *et al*., 2020b; Gong *et al*., 2021). However, the composition of resistant populations is variable and appears to differ between survey years and across regions. Field surveys of *S. avenae* in Ireland indicate that the composition of individuals containing the *kdr*-SR heterozygous mutation can range from 25-54% (Walsh *et al*., 2020b). Resistance against pyrethroids was recently reported in an *R. padi* population collected from Jingyang, Shaaxi Province, China (Wang *et al*., 2020) and subsequent field surveys have detected additional pyrethroid-resistant populations in multiple locations across China (Gong *et al*., 2021). According to the Arthropod Pesticide Resistance Database, a global databank of insecticide resistance cases, no other occurrences of pyrethroid resistance in *R. padi* have been reported. This indicates that full pyrethroid resistance is yet to evolve, or be detected, in *R. padi* populations outside of China.

The development of insecticide resistance can be monitored through dose-response assays to detect the emergence of populations showing reduced sensitivity to insecticides. *R. padi* populations with reduced sensitivity to pyrethroids have been recently detected in Ireland (Walsh *et al*., 2020a) and Australia (Umina *et al*., 2020), suggesting that resistance could be evolving. However, on average, field concentrations of pyrethroids are still effective at controlling over 90% of the aphid population (Zuo *et al*., 2016; Umina *et al*., 2020; Walsh *et al*., 2020a).

The lack of high prevalence of resistant populations across regions and years (Walsh *et al*., 2020b; Gong *et al*., 2021) suggests that fitness consequences could be associated with insecticide resistance traits. Recent research has provided some evidence to support this: it was recently reported that *S. avenae* populations with heterozygous *kdr*-SR resistance to pyrethroids exhibit increased vulnerability to the parasitoid *Aphidius ervi* (Jackson *et al*., 2020). Studies have also identified additional fitness trade-offs resulting from *kdr*-SR heterozygous resistance to pyrethroids, including lower aphid abundance and reduced growth in *S. avenae* populations (Jackson *et al*., 2020) and reduced fecundity in *R. padi* populations carrying *super-kdr* resistance (Wang *et al*., 2021).

A key driver of phenotypic diversity in aphid populations is the presence of facultative endosymbionts (Zytynska & Weisser, 2016; Zytynska *et al*., 2021). The majority of aphid species form an essential relationship with the endosymbiont *Buchnera aphidicola*, with *B. aphidicola* providing nutritional supplementation to the aphid diet (Sasaki *et al*., 1991; Douglas & Prosser, 1992). Aphids can also form non-essential, or facultative, relationships with a range of additional endosymbionts (Zytynska & Weisser, 2016; Guo *et al*., 2017). The most common facultative endosymbionts detected in aphid populations are *Spiroplasma spp., Regiella insecticola, Hamiltonella defensa, Rickettsiella sp., Fukatsia symbiotica* (previously pea aphid x-type symbiont, *PAXS), Seratia symbiotica, Rickettsia spp*., and *Arsenophonus spp*. (Zytynska & Weisser, 2016; Guo *et al*., 2017; Beekman *et al*., 2022). These facultative endosymbiotic relationships occur naturally in aphid populations (Henry *et al*., 2015; Guo *et al*., 2019; Leybourne *et al*., 2020a; Beekman *et al*., 2022). The phenotypic consequence of endosymbiont infection is not always clear, and the phenotypic traits conferred by a specific endosymbiont species are not consistently observed between aphid species, aphid genotypes within the same species, or even between different endosymbiont strains (Vorburger *et al*., 2010; Cayetano *et al*., 2015; McLean & Godfray, 2015; Oliver & Higashi, 2019). One common beneficial trait that is often conferred through facultative endosymbiont infection across a range of aphid-endosymbiont combinations is resistance against parasitoid wasps (Oliver *et al*., 2003, 2009; Asplen *et al*., 2014; Leybourne *et al*., 2020a). A diverse range of other phenotypic traits that can be conferred by endosymbiont infection have also been described, including lower fecundity (Zytynska *et al*., 2021).

In cereal aphids, endosymbiont-conferred phenotypes include protection against parasitoid wasps (Leybourne *et al*., 2020a), altered feeding behaviour (Leybourne *et al*., 2020b), adjusted life-history parameters, including reduced growth and development (Liu *et al*., 2019; Leybourne *et al*., 2020a; Luo *et al*., 2020), and moderate increase in susceptibility to bacterial pathogens (Álvarez-Lagazzi *et al*., 2021). Recently, studies have indicated that endosymbiont infection can also influence the susceptibility of the aphid host to insecticides. A study examining aphid susceptibility to a range of insecticides found that wheat aphids, *S. miscanthi*, infected with *H. defensa* were more susceptible to low concentrations of insecticide when compared with uninfected aphids (Li *et al*., 2021). This indicates that there could be a phenotypic link between facultative endosymbiont communities and aphid susceptibility, or tolerance, to insecticides. An association between endosymbiont infection and insecticide resistance could provide an explanation for high variation in endosymbiont prevalence and *kdr*-SR prevalence in aphid populations (Guo *et al*., 2019; Walsh *et al*., 2020b).

Here, we report the results of pyrethroid dose-response bio-assays for cereal aphid populations sampled from a key cereal production region in Northern Germany. We sampled 25 *S. avenae* and seven *R. padi* populations from 13 field sites. We find that, for *S. avenae*, the level of insecticide susceptibility is highly variable, with populations sensitive and tolerant to pyrethroid exposure, including populations collected from the same field. In *R. padi* populations, we find evidence for decreased sensitivity, indicating that resistance to pyrethroids is starting to evolve in German *R. padi* populations. Furthermore, we explore the hypothesis that endosymbiont infection increases aphid susceptibility to insecticides by characterising the endosymbiont communities of these populations.

## Methods

### Aphid populations and sampling

Cereal aphid populations were sampled in summer and autumn 2021 from 13 agricultural fields in Lower Saxony, Germany (Fig. 1). Sample sites comprised 12 winter cereal fields (collected in summer) and one winter rapeseed field (collected in autumn). Single adult aphids (apterous or alate) were collected and used to establish laboratory populations; from the 13 fields 25 *Sitobion avenae* and seven *Rhopalosiphum padi* populations were established. Populations were maintained on one-week old wheat plants (approximately GS 11-13) in ventilated plastic cups under glasshouse conditions. Where multiple samples were collected from one field the samples were either collected on different dates or with a 20 m minimum distance between sampling points.

**Fig. 1:**
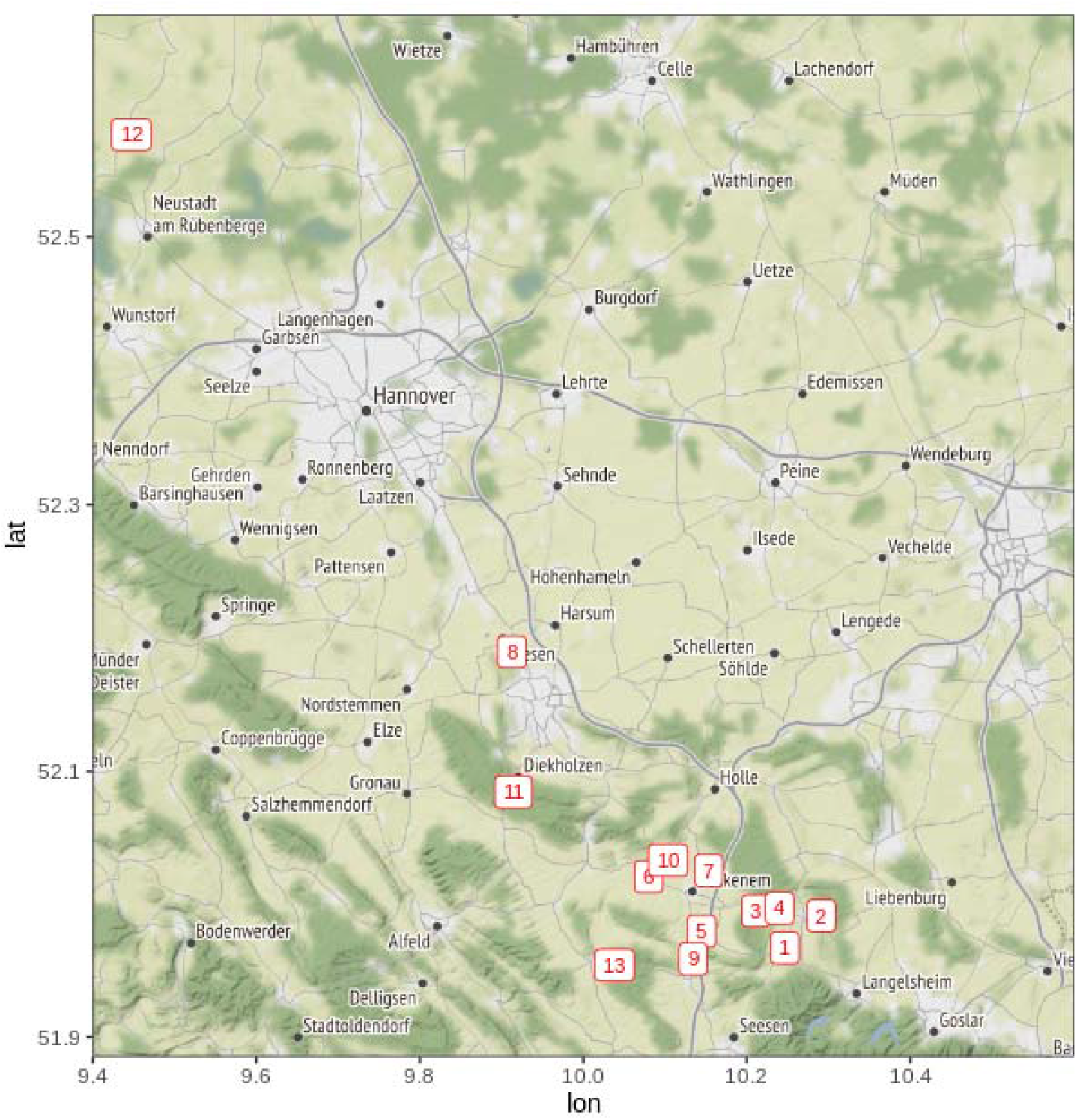
Location of the 13 sample sites. Field 12 was a winter rapeseed field, all other fields were winter wheat

### Insecticide sensitivity testing

Aphid populations were screened for susceptibility and sensitivity to the synthetic pyrethroid Decis Forte^®^ (Bayer CropScience, Germany), a Class 3A synthetic pyrethroid (active ingredient deltamethrin at 100 g L^-1^ formulation). This insecticide was selected as it is approved for use on arable and field crops in Germany. A stock solution was prepared in water at a concentration comparable to the recommended field rate, equating to a concentration of 357 mg a.i. L^-1^. Serial dilutions were prepared from the stock solution. Five insecticide dilutions were used in the assay: stock, 10^-1^, 10^-2^, 10^-3^, 10^-4^, with distilled water included as a negative control.

The insecticide sensitivity assays broadly followed the IRAC leaf-dip method (IRAC, 2016) and the method deployed by (Umina *et al*., 2020). Briefly, *c*. 25 mm sections of wheat leaves were submerged for *c. 10* s in one of the test solutions. Control leaves were dipped first, then the leaves were dipped sequentially from the lowest concentration (10^-4^) to the field rate stock solution. Once dipped, leaves were left to dry on paper towels for approximately 1 h before they were placed abaxial side up on agar (1 g L^-1^) in a plastic Petri Dish; a droplet of water was added to the surface of the agar to aid leaf adhesion. Aphids were transferred to each Petri Dish using a fine-haired paintbrush, Petri Dishes were moved to a controlled environment room (20°C ± 2°C, L16:D8), and Petri Dishes were inverted to simulate aphid feeding from the underside of the leaf. Between 4 – 8 aphids were transferred to each Petri Dish. After 48 h aphids were scored as either alive, moribund, or dead. Aphids were classed as alive if they were able to return to an upright position when placed on their back (i.e., they were capable of coordinated movement). Moribund and dead aphids were grouped together as “affected”, in-line with previous dose-response assays (Foster *et al*., 2012; Umina *et al*., 2020).

### DNA extraction and diagnostic PCR for endosymbiont characterisation

A sample of five aphids (mixture of apterous adults and nymphs) were collected from each population and DNA was extracted using the Norgen® Plant and Fungi DNA extraction kit (Norgen Biotek, Germany) following manufacturer’s instructions. An extraction blank was included with each batch of extractions.

Successful DNA extraction was confirmed using a PCR marker for the primary aphid symbiont *Buchnera aphidocola*. The presence of facultative endosymbionts was determined using a three-step multiplex diagnostic PCR assay (Beekman *et al*., 2022). Multiplex assays were used to detect the presence of the main aphid secondary endosymbionts: *Spiroplasma spp., Regiella insecticola, Hamiltonella defensa, Rickettsiella sp., Fukatsuia symbiotica, Seratia symbiotica, Rickettsia spp*., and *Arsenophonus spp*. All PCR primer details are described in Table S1. PCR assays were conducted in a final reaction volume of 12 μL consisting of: 2 μL DNA, 6 μL 2X Kappa2G Fast PCR Ready Mix (Merck, Germany). Primer concentrations and volumes differed between the multiplex assays and are detailed in Table S1. The final reaction mixture was made to 12 μL using nuclease-free DEPC-treated water (CarlRoth, Germany). PCR conditions followed Beekman *et al*., 2022, i.e.: denaturation at 94°C for 3 min followed by 35 cycles of 94°C for 30 s, 58°C for 30 s, and 72°C for 60 s with a final extension step at 72°C for 10 min. Positive DNA (mixed DNA containing positive DNA extracts for all target endosymbionts) was included as a positive control, an extraction blank was used as an extraction negative control, and DNA-free PCR mastermix was included as PCR negative control. Endosymbiont presence was detected by separation of PCR products on a 1% agarose gel stained with GelRed® (Biotium, Germany), and reactions were visualised under UV light; a 100 bp DNA ladder (ThermoFischer, Germany) was used to estimate band size. Positive identification of the presence of endosymbionts in the multiplex assay were confirmed in additional singleplex assays. All PCR assays were conducted in a Biometra TRIO 48, Thermocycler (Analytik Jena, Germany).

### Statistical analysis

All statistical analysis was carried out using R (v.4.1.2) and R Studio (v.1.3.1093). The estimated concentration of active ingredient required to achieve 50% mortality (EC_50_ value), EC_50_ 95% confidence intervals, slope, intercept, and associated standard errors for the dose response curve were calculated for each aphid population using probit estimation regression (Finney, 1952). To achieve this, the “ProbitEPA” function in the R package ecotoxicology (v.1.0.1) was used. Differences in the dose response between populations was detected using an ANOVA, as done in similar studies (Umina *et al*., 2020); to achieve this the intercepts of the dose response curves were estimated using linear models with aphid population, facultative endosymbiont infection status, and facultative endosymbiont diversity (Simpson’s diversity) included as explanatory variables in individual models. Linear models were tested for significance using Type-II ANOVA. Simpson’s diversity was calculated using the vegan package (v.2.5-7). Where significant differences in model intercepts were detected, the differing aphid populations were identified by observing the overlapping confidence intervals. This method has been used previously to identify differences in insecticide susceptibility between aphid populations from estimated concentration of active ingredient required to achieve 50% population mortality (EC_50_) values (Foster *et al*., 2012).

## Results

### *Pyrethroid sensitivity is variable in* Sitobion avenae *populations*

Based on mortality at field rate concentration (i.e., mortality in the stock treatment, 357 mg a.i. L^-1^ deltamethrin), *S. avenae* populations were grouped into four broad categories (Table 1): Susceptible (mortality at field concentration 95-100%), moderately susceptible (mortality at field concentration 90-94%), moderately tolerant (mortality at field concentration 71-89%), and tolerant (mortality at field concentration ≤ 70%). Field rate concentration only achieved complete aphid control in five of the 25 *S. avenae* populations, namely populations SA3, SA-5, SA-6, SA-11, and SA-23 (Fig. 2). Three *S. avenae* populations (SA-13, SA-15, SA-16) showed tolerance to pyrethroid exposure, with ≤ 70% population mortality following exposure to field rate concentrations of deltamethrin (Fig. 2; Table 1).

**Table 1:** Insecticide bio-assay results

| Clone name | Species | Field | Week sampled | Number tested | Slope coefficient (±SE) | EC<sub>50</sub> (mg a.i. L<sup>-1</sup>) | 95% confidence interval | Resistance Category |
| --- | --- | --- | --- | --- | --- | --- | --- | --- |
| SA-1<sup>AB</sup> | *S. avenae* | 6 | 21.06.2021 | 153 | 0.50 (0.08) | 0.62 | 0.09 – 3.00 | Susceptible |
| SA-2<sup>BCDEF</sup> | *S. avenae* | 5 | 21.06.2021 | 173 | 0.70 (0.18) | 9.59 | 2.99 – 36.71 | Susceptible |
| SA-3<sup>AB</sup> | *S. avenae* | 1 | 21.06.2021 | 126 | 0.89 (0.11) | 1.81 | 0.49 – 6.11 | Susceptible |
| SA-4<sup>ABCD</sup> | *S. avenae* | 7 | 21.06.2021 | 123 | 0.67 (0.11) | 1.91 | 0.37 – 8.57 | Susceptible |
| SA-5<sup>A</sup> | *S. avenae* | 10 | 21.06.2021 | 168 | 0.79 (0.09) | 0.38 | 0.10 – 1.13 | Susceptible |
| SA-6<sup>AB</sup> | *S. avenae* | 2 | 21.06.2021 | 174 | 0.66 (0.18) | 1.09 | 0.28 – 3.64 | Susceptible |
| SA-8<sup>CDEF</sup> | *S. avenae* | 5 | 21.06.2021 | 124 | 0.69 (0.15) | 15.67 | 3.84 – 96.17 | Moderately Susceptible |
| SA-9<sup>CDEF</sup> | *S. avenae* | 1 | 21.06.2021 | 126 | 0.57 (0.38) | 24.62 | 5.03 – 295.95 | Moderately Tolerant |
| SA-10<sup>CDEF</sup> | *S. avenae* | 1 | 21.06.2021 | 176 | 0.60 (0.33) | 20.39 | 5.64 – 114.40 | Moderately Tolerant |
| SA-11<sup>BCDE</sup> | *S. avenae* | 7 | 21.06.2021 | 192 | 1.05 (0.18) | 4.14 | 1.17 – 10.04 | Susceptible |
| SA-12<sup>BCDEF</sup> | *S. avenae* | 3 | 21.06.2021 | 192 | 0.51 (0.28) | 8.26 | 2.09 – 43.62 | Moderately Susceptible |
| SA-13<sup>EF</sup> | *S. avenae* | 6 | 21.06.2021 | 174 | 0.38 (0.09) | 68.56 | 10.19 – 3319.38 | Tolerant |
| SA-14<sup>CDEF</sup> | *S. avenae* | 6 | 21.06.2021 | 192 | 0.48 (0.12) | 24.52 | 5.75 – 215.58 | Moderately Tolerant |
| SA-15<sup>F</sup> | *S. avenae* | 7 | 21.06.2021 | 144 | 0.38 (0.10) | 190.08 | 19.80 – 11356.90 | Tolerant |
| SA-16<sup>EF</sup> | *S. avenae* | 7 | 21.06.2021 | 156 | 0.45 (0.11) | 80.53 | 13.31 – 3572.30 | Tolerant |
| SA-17<sup>CDEF</sup> | *S. avenae* | 9 | 21.06.2021 | 126 | 0.75 (0.16) | 20.29 | 5.39 – 114.46 | Susceptible |
| SA-18<sup>ABCDE</sup> | *S. avenae* | 3 | 21.06.2021 | 156 | 0.69 (0.08) | 2.86 | 0.77 – 10.34 | Moderately Susceptible |
| SA-19<sup>BCDEF</sup> | *S. avenae* | 7 | 21.06.2021 | 174 | 0.63 (0.30) | 9.08 | 2.60 – 39.21 | Moderately Susceptible |
| SA-20<sup>CDEF</sup> | *S. avenae* | 3 | 21.06.2021 | 174 | 0.53 (0.17) | 23.01 | 5.57 – 184.19 | Moderately Tolerant |
| SA-21<sup>DEF</sup> | *S. avenae* | 3 | 05.07.2021 | 126 | 0.59 (0.13) | 39.30 | 8.27 – 573.09 | Moderately Tolerant |
| SA-22<sup>CDEF</sup> | *S. avenae* | 9 | 21.06.2021 | 174 | 0.52 (0.17) | 16.87 | 4.07 – 120.94 | Moderately Tolerant |
| SA-23<sup>AB</sup> | *S. avenae* | 13 | 21.06.2021 | 149 | 0.62 (0.18) | 0.85 | 0.16 – 3.33 | Susceptible |
| SA-24<sup>DEF</sup> | *S. avenae* | 6 | 05.07.2021 | 173 | 0.63 (0.19) | 22.71 | 6.49 – 123.98 | Moderately Susceptible |
| SA-25<sup>EF</sup> | *S. avenae* | 9 | 21.06.2021 | 174 | 0.63 (0.64) | 33.94 | 9.74 – 209.92 | Moderately Tolerant |
| SA-26<sup>BCDEF</sup> | *S. avenae* | 12 | 04.10.2021 | 170 | 0.63 (0.13) | 9.15 | 2.57 – 41.39 | Susceptible |
| RP-1<sup>ZY</sup> | *R. padi* | 2 | 21.06.2021 | 189 | 0.81 (0.07) | 1.13 | 0.37 – 3.11 | Susceptible |
| RP-2<sup>ZY</sup> | *R. padi* | 5 | 05.07.2021 | 192 | 0.73 (0.10) | 1.55 | 0.49 – 4.54 | Susceptible |
| RP-3<sup>Z</sup> | *R. padi* | 5 | 21.06.2021 | 189 | 0.66 (0.15) | 0.44 | 0.11 – 1.40 | Susceptible |
| RP-4<sup>ZY</sup> | *R. padi* | 11 | 21.06.2021 | 192 | 0.63 (0.18) | 1.41 | 0.39 – 4.63 | Susceptible |
| RP-5<sup>Y</sup> | *R. padi* | 13 | 21.06.2021 | 192 | 1.50 (0.11) | 5.32 | 2.51 – 11.29 | Susceptible |
| RP-6<sup>ZY</sup> | *R. padi* | 4 | 21.06.2021 | 187 | 0.82 (0.09) | 1.63 | 0.55 – 4.55 | Susceptible |
| RP-7<sup>ZY</sup> | *R. padi* | 4 | 05.07.2021 | 192 | 0.78 (0.07) | 2.03 | 0.69 – 5.69 | Susceptible |

The EC<sub>50</sub> values for populations followed by the same letter do not differ significantly, based on observation of overlapping confidence intervals (Foster *et al.*, 2011). Populations have been allocated a resistance category based on the observed mortality (% aphids affected) at the field concentration (375 mg a.i. / L) treatment: Susceptible (95-100% mortality), moderately susceptible (90-94% mortality), moderately tolerant (71-89% mortality), tolerant (≤ 70% mortality).

**Fig. 2:**
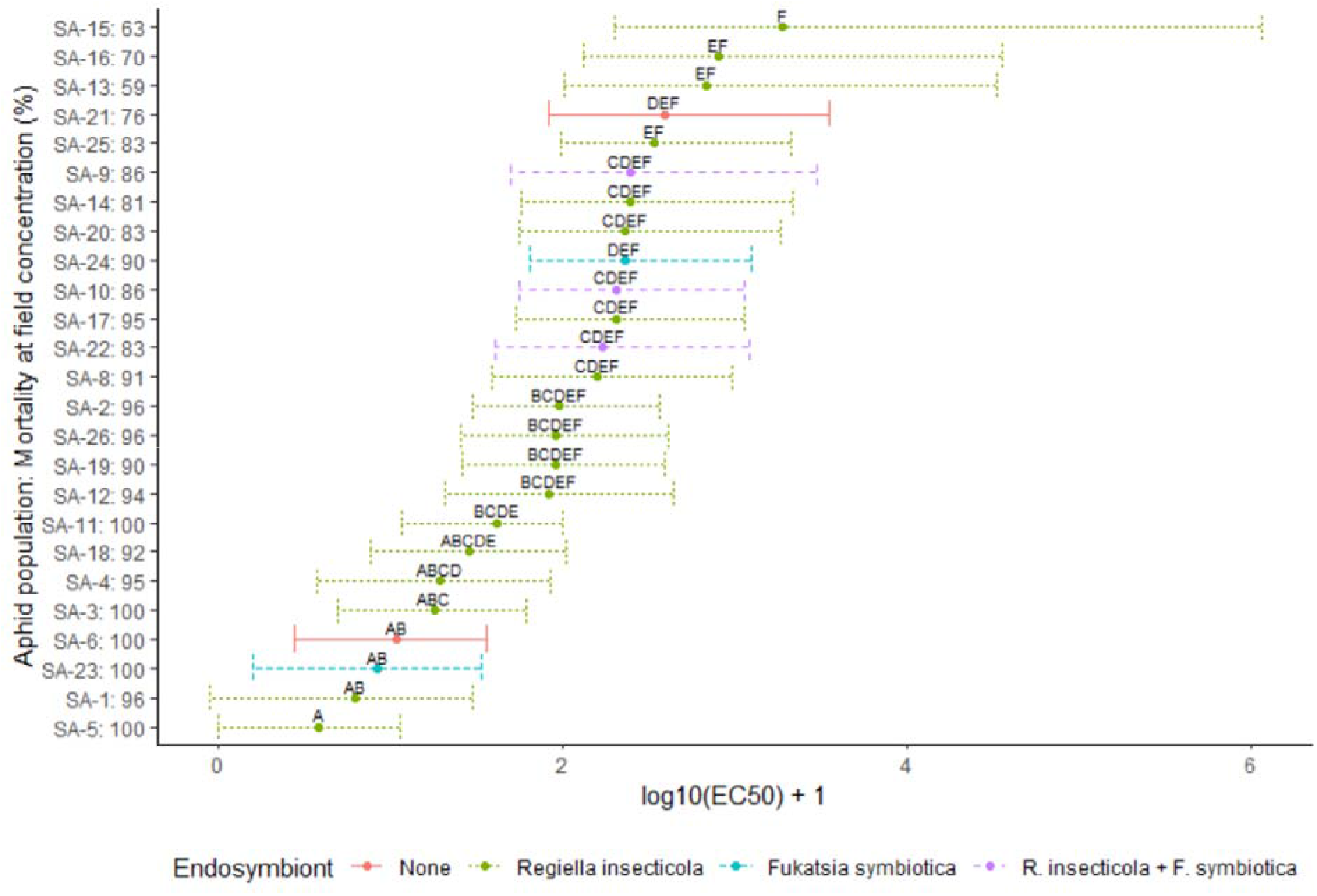
Estimated EC_50_ values and 95% confidence intervals for *Sitobion avenae* populations. EC_50_ values are shown on the log_10_+1 scale to aid interpretation. Aphid population on the y-axis is followed by the percentage population control at field rate concentration. Letters indicate differences between the aphid populations based on overlapping confidence intervals.

The estimated effective dose required for 50% population control (EC_50_) ranged from 0.38 mg a.i. L^-1^ to 190.08 mg a.i. L^-1^ deltamethrin (Table 1). Comparison of the dose response model intercepts highlighted differences in EC_50_ amongst the *S. avenae* populations examined (F_24,92_ = 4.16; p = <0.001). Observation of the overlapping 95% confidence intervals (Fig. 2; Table 1) indicates that differences in EC_50_ are between the three tolerant populations (SA-13, SA-15, SA-16) and seven of the ten susceptible populations (SA-1, SA-3, SA-4, SA-5, SA-6, SA-11, SA-23).

### *Decreased pyrethroid sensitivity in a* Rhopalosiphum padi *population collected from Germany*

Based on mortality at field rate concentration, all *R. padi* populations were categorised as susceptible to deltamethrin (mortality >95%; Table 2; Fig. 3); however, two populations (RP-5 and RP-6) had a reduced mortality of 96% (Fig. 3).

**Table 2:** Endosymbiont profiles of the 32 aphid populations

| Clone name | Species | *B.a*<br>(primary) | *Spi* | *R.i.* | *H.d.* | *R-siella* | *F. s.* | *S.s.* | *R-tsia* | *Ars.* |
| --- | --- | --- | --- | --- | --- | --- | --- | --- | --- | --- |
| SA-1 | *S. avenae* | + | - | + | - | - | - | - | - | - |
| SA-2 | *S. avenae* | + | - | + | - | - | - | - | - | - |
| SA-3 | *S. avenae* | + | - | + | - | - | - | - | - | - |
| SA-4 | *S. avenae* | + | - | + | - | - | - | - | - | - |
| SA-5 | *S. avenae* | + | - | + | - | - | - | - | - | - |
| SA-6 | *S. avenae* | + | - | - | - | - | - | - | - | - |
| SA-8 | *S. avenae* | + | - | + | - | - | - | - | - | - |
| SA-9 | *S. avenae* | + | - | + | - | - | + | - | - | - |
| SA-10 | *S. avenae* | + | - | + | - | - | + | - | - | - |
| SA-11 | *S. avenae* | + | - | + | - | - | - | - | - | - |
| SA-12 | *S. avenae* | + | - | + | - | - | - | - | - | - |
| SA-13 | *S. avenae* | + | - | + | - | - | - | - | - | - |
| SA-14 | *S. avenae* | + | - | + | - | - | - | - | - | - |
| SA-15 | *S. avenae* | + | - | + | - | - | - | - | - | - |
| SA-16 | *S. avenae* | + | - | + | - | - | - | - | - | - |
| SA-17 | *S. avenae* | + | - | + | - | - | - | - | - | - |
| SA-18 | *S. avenae* | + | - | + | - | - | - | - | - | - |
| SA-19 | *S. avenae* | + | - | + | - | - | - | - | - | - |
| SA-20 | *S. avenae* | + | - | + | - | - | - | - | - | - |
| SA-21 | *S. avenae* | + | - | - | - | - | - | - | - | - |
| SA-22 | *S. avenae* | + | - | + | - | - | + | - | - | - |
| SA-23 | *S. avenae* | + | - | - | - | - | + | - | - | - |
| SA-24 | *S. avenae* | + | - | - | - | - | + | - | - | - |
| SA-25 | *S. avenae* | + | - | + | - | - | - | - | - | - |
| SA-26 | *S. avenae* | + | - | + | - | - | - | - | - | - |
| RP-1 | *R. padi* | + | - | - | - | - | - | - | - | - |
| RP-2 | *R. padi* | + | - | - | + | - | - | - | - | - |
| RP-3 | *R. padi* | + | - | - | + | - | - | - | - | - |
| RP-4 | *R. padi* | + | - | - | - | - | - | - | - | - |
| RP-5 | *R. padi* | + | - | - | - | - | - | - | - | - |
| RP-6 | *R. padi* | + | - | - | - | - | + | - | - | - |
| RP-7 | *R. padi* | + | - | - | - | - | + | - | - | - |

Symbiont abbreviations: *B.a* (*B. aphidicola*; essential primary endosymbiont), *Spi* (*Spiroplasma* spp.), *R.i.* (*Regiella insecticola*), *H.d.* (*Hamiltonella defensa*), *R-siella* (*Rickettsiella* sp.), *F.s.* (*Fukatsia symbiotica*), *S.s.* (*Seratia symbiotica*), *R-tsia* (*Rickettsia* spp.), and *Ars.* (*Arsenophonus* spp).

**Fig. 3:**
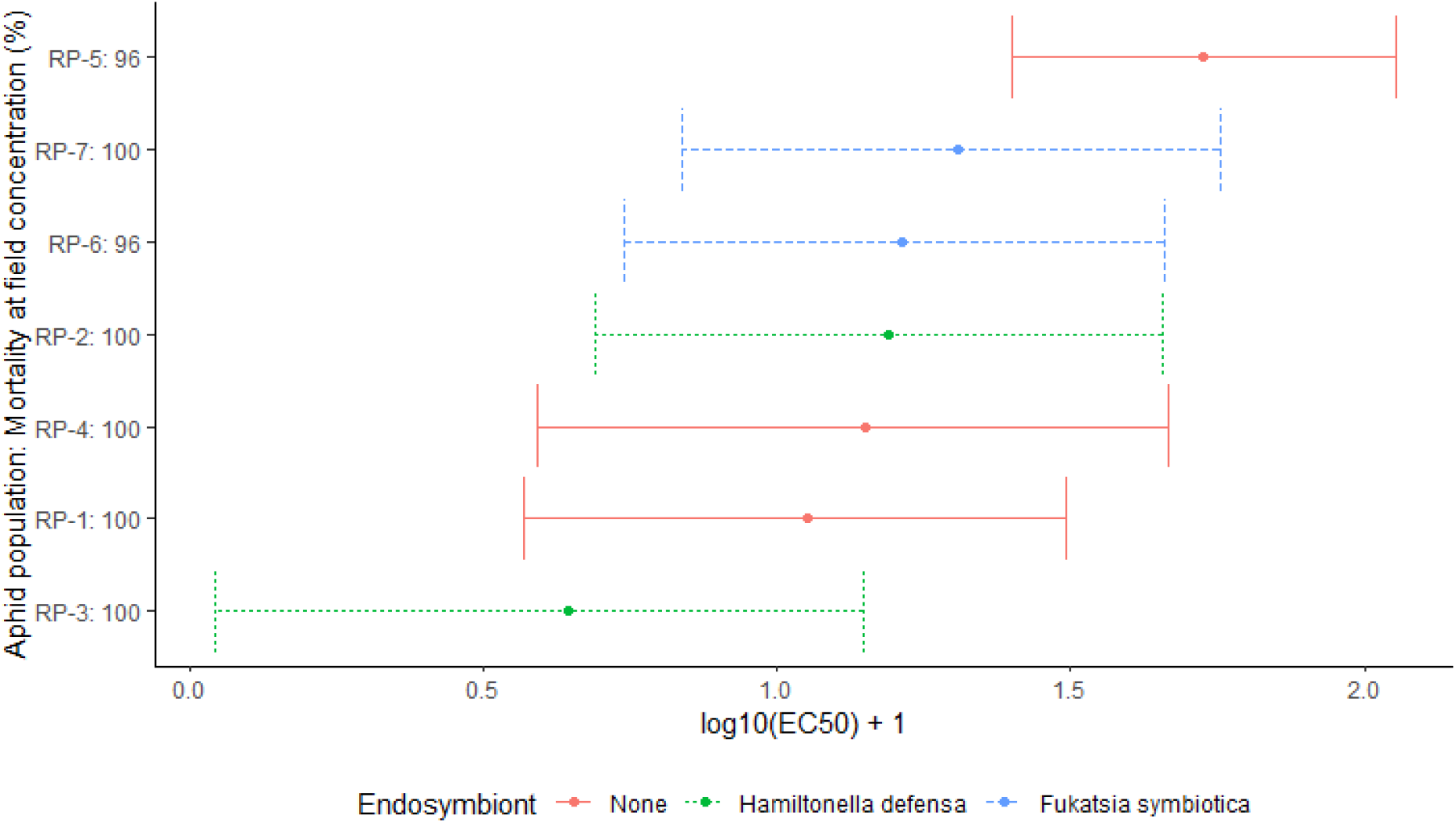
Estimated EC50 values and 95% CI for *Rhopalosiphum padi* populations. EC_50_ values are shown on the log_10_+1 scale to aid interpretation. Aphid population on the y-axis is followed by the percentage population control at field rate concentration.

The estimated effective dose required for 50% population control (EC_50_) ranged from 0.44 mg a.i. L^-1^ to 5.32 mg a.i. L^-1^ deltamethrin (Table 1). Comparison of model intercepts highlighted differences in EC_50_ amongst the *R. padi* populations examined (F_6,27_ = 7.43; p = <0.001). Observation of the overlapping 95% confidence intervals (Fig. 3; Table 1) indicates that differences in EC_50_ are between one of the populations with reduced mortality, RP-5, and the susceptible population RP-3 (Fig. 3; Table 1).

### Facultative endosymbionts occur at high frequencies in aphid populations but they do not influence pyrethroid sensitivity

Of the 32 aphid populations 27 were infected with at least one facultative endosymbiont (Table 2). The endosymbiont community differed between the two cereal aphid species: endosymbionts were detected in 92% of *S. avenae* populations and 57% of *R. padi* populations (Table 2). In *S. avenae* endosymbiont communities were dominated by *R. insecticola*, with *R. insecticola* present in 72% of *S. avenae* populations. Low levels of infection with *F. symbiotica* (8%) and co-infection of *R. insecticola* and *F. symbiotica* (12%) were detected in the *S. avenae* populations. For *R. padi*, the defensive endosymbiont *H. defensa* was detected in 28% of the populations and *F. symbiotica* was detected in 28% of the populations. For both aphid species, neither endosymbiont infection status nor endosymbiont diversity influenced aphid sensitivity to deltamethrin: *S. avenae* facultative endosymbiont infection status (F_1,23_ = 0.37; p = 0.778; Fig. 2) and diversity (F_1,23_ = 0.06; p = 0.816), *R. padi* facultative endosymbiont infection status (F_2,4_ = 0.65; p = 0.569; Fig. 3) and diversity F_2,4_ = 1.11; p = 0.341).

## Discussion

Our results provide insecticide dose-response data for 32 aphid populations against a synthetic pyrethroid approved for aphid management in arable crops in Germany. We observe wide variation in dose-response amongst the 25 *S. avenae* populations and we detect reduced sensitivity to deltamethrin in one of the *R. padi* populations tested. We detect natural infection with the facultative endosymbionts *R. insecticola* in *S. avenae, H. defensa* in *R. padi*, and *F symbiotica* in both species, including co-infection of *R. insecticola* and *F. symbiotica* in a subset of *S. avenae* populations.

For the *S. avenae* populations we detected variable response to pyrethroid exposure, with EC_50_ values ranging from 0.38 mg a.i. L^-1^ to 190.08 mg a.i. L^-^. This wide variation in dose response in *S. avenae* indicates that our *S. avenae* populations comprise individuals that are highly sensitive to pyrethroids, tolerant to pyrethroids, and populations with intermediate susceptibilities. Indeed, the low EC_50_ value of 0.38 mg a.i. L^-1^ detected in SA-5 is comparable with EC_50_ values in *S. avenae* populations that are sensitive to pyrethroid exposure (0.50 mg Bifenthrin L^-1^; 2.40 mg Beta-cypermethrin L^-1^; Gong *et al*., 2021) and the high EC_50_ values estimated for the three tolerant populations, SA-15 (190.08 mg a.i. L^-1^), SA-16 (80.53 mg a.i. L^-1^), SA-13 (68.56 mg a.i. L^-1^), are similar to the EC_50_ values reported for *S. avenae* populations tolerant to pyrethroids, including those that harbour heterozygous *kdr*-SR resistance (Foster *et al*., 2014; Walsh *et al*., 2020a; Gong *et al*., 2021). Our field survey also indicates that pyrethroid susceptibility is highly variable between populations collected from the same field. We sampled five *S. avenae* populations from field 7 and the estimated EC_50_ for these populations ranged from 1.91 mg a.i. L^-1^ 190.08 mg a.i. L^-1^. Based on the observed population control at field rate concentrations, the aphids collected from this field contained populations grouped into the susceptible, moderately tolerant, and tolerant categories. Similar observations were made for other locations where more than one population was sampled: Field 3 contained four *S. avenae* populations comprising those categorised as moderately tolerant and moderately susceptible; field 9 had three populations comprising those categorised as susceptible and moderately tolerant; four individuals were collected from field 6 comprising those categorised as susceptible, moderately tolerant, and tolerant; and three populations were sampled from field 1 comprising populations categorised into susceptible and moderately tolerant. This diverse range of susceptibility and tolerance to pyrethroids, including wide variation within the same sampling locations, is in-line with recent surveys conducted in Ireland (Walsh *et al*., 2020b) and China (Gong *et al*., 2021). However, these studies did not relate insecticide susceptibility or tolerance to the presence or absence of facultative endosymbionts within the local aphid populations.

In our *R. padi* populations we detected reduced sensitivity to the pyrethroid deltamethrin in one population. This difference is between the most sensitive aphid population, RP-3 estimated EC_50_ 0.44 mg a.i. L^-1^, and population RP-5, estimated to have an increased EC_50_ of 5.32 mg a.i. L^-1^. We also detected a reduction in the level of population control in RP-5, with only 96% of the aphid population effectively managed at the field rate concentration of 357 mg a.i. L ^-1^, compared with 100% population control in RP-3. Our increased EC_50_ for RP-5 represents the first observation of reduced sensitivity to pyrethroids in *R. padi* populations collected in Germany. Reduced sensitivity against pyrethroids has recently been reported in three *R. padi* populations in Australia, these three populations were estimated to have EC_50_ values of 15.34, 16.22, and 24.57 mg a.i. L^-1^ (Umina *et al*., 2020). These values are at least three-fold higher than the EC_50_ value estimated for RP-5 and are associated with a further reduction in effective population control under field rate concentrations, down to 91% effective control (Umina *et al*., 2020), compared with 96% control in RP-5. Recent surveys in Ireland have also detected reduced pyrethroid sensitivity in one *R. padi* population, where EC_50_ levels were compared with a susceptible *kdr*-SS *S. avenae* population (Walsh *et al*., 2020a). Although we detect reduced sensitivity in *R. padi* in Germany, effective population control remains relatively high at 96%, thus this should not adversely affect aphid management strategies or impact crop yields in the immediate term. However, this finding indicates that resistance to pyrethroids is starting to evolve and, upon the evolution of *kdr*-SR resistance, the impact on crop yields could become more apparent as conventional control strategies fail. Surveys should be continued in order to monitor the development of the situation over the coming years while developing and prioritising the use of alternative strategies for aphid management that do not rely on resistance-inducing insecticides. In China, decreased sensitivity to pyrethroids was first detected in populations sampled in 2013 from multiple locations across China, including Xianyang, Shaaxi Province (Zuo *et al*., 2016). *Super-kdr*-SR heterozygous resistance against pyrethroids (shown to be effective against two active ingredients: alpha-cypermethrin and deltamethrin) was detected six years later from populations collected in 2019 from the same Province (Wang *et al*., 2020).

Aphids are often considered as a single homogeneous population, however it is clear that aphids can comprise populations with contrasting intra-species diversity. Intra-species diversity in cereal aphids is associated with genetic diversity and the composition of endosymbiont communities (Alkhedir *et al*., 2013; Malloch *et al*., 2016; Guo *et al*., 2019; Leybourne *et al*., 2020a). This intra-species diversity can affect the aphid phenotype, and there is evidence that this could include heightened aphid susceptibility to insecticides (Li *et al*., 2021). In order to examine whether natural occurrence of facultative endosymbionts influences aphid sensitivity to pyrethroids, we characterised the facultative endosymbiont community of the 32 aphid populations and related this to the results of the dose-response assays. We detected facultative endosymbionts in 92% of the *S. avenae* populations and 57% of the *R. padi* populations. These natural levels of endosymbiont infection are similar to those reported in previous endosymbiont surveys (Alkhedir *et al*., 2013; Łukasik *et al*., 2013; Fakhour *et al*., 2018; Guo *et al*., 2019). Our *S. avenae* populations showed a high prevalence of *R. insecticola* infection (72%). This is comparable with infection levels detected in Morocco, 75% *S. avenae* population infection with *R. insecticola* (Fakhour *et al*., 2018), and above levels previously observed in *S. avenae* sampled from Germany, 50% *R. insecticola* infection (Alkhedir *et al*., 2013). Similarly, *H. defensa* usually occurs at low-to-moderate frequency in *R. padi* populations, with previous studies reporting infection frequencies between 10-38% (Guo *et al*., 2019; Leybourne *et al*., 2020a), comparable with 29% of *R. padi* populations detected to be infected with *H. defensa* in our aphid populations. We detected *F. symbiotica* in a small proportion of our *S. avenae* and *R. padi* populations. *F. symbiotica* can occur at high levels in *A. pisum* populations (Zytynska & Weisser, 2016) but rarely infects other aphid species, occurring at low frequencies where it is detected (Łukasik *et al*., 2013; Zytynska & Weisser, 2016).

Although we detected variation in endosymbiont infection frequencies across our populations, with 84% of populations carrying at least one facultative endosymbiont and 9% carrying two facultative endosymbiont species, we did not detect any link between endosymbiont infection and heightened susceptibility to insecticides. This is in contrast with recent lab studies, where an association between endosymbiont infection and heightened insecticide susceptibility has been reported (Skaljac *et al*., 2018; Li *et al*., 2021). Recent research has shown that artificial inoculation with *H. defensa* in the grain aphid *S. miscanthi* increases aphid sensitivity to a range of insecticides at low concentrations, including neonicotinoids and diamides (Li *et al*., 2021). Similar observations have been made in pea aphids (*Acyrthosiphon pisum*) infected with the endosymbiont *S. symbiotica*, where symbiont-infected aphids were more susceptible to low concentrations of several insecticides, including carbamates, neonicotonoids, tetronic and tertamic acid derivatives, and diamides (Skaljac *et al*., 2018). These studies also show that the EC_50_ values are lower for symbiont-infected aphids compared with aphid populations that do not contain facultative endosymbiont communities (Skaljac *et al*., 2018; Li *et al*., 2021). Although these studies did not examine the relationship between endosymbiont presence and susceptibility to pyrethroids, they still showcase a link between endosymbiont infection and heightened susceptibility to insecticide exposure in aphid populations. However, it should be noted that these were artificially manipulated populations developed through the endosymbiont removal and infection to establish desired endosymbiont communities under lab conditions, not comparisons of natural infections (Skaljac *et al*., 2018; Li *et al*., 2021). The next stage of research would be to examine this association under field conditions, and to examine this across a broader range of insecticides, including the important pyrethroids.

One caveat of our study was our lack of a characterised *kdr*-SS homozygous pyrethroid susceptible reference clone. A susceptible clone can be used as a reference baseline in order to calculate resistance ratios for each tested population and to act as an internal reference (Walsh *et al*., 2020a; Wang *et al*., 2020), although this is not included in every survey (Umina *et al*., 2020; Gong *et al*., 2021). The calculated EC_50_ values for our most highly sensitive aphid population for each species, *R. padi* (0.44 mg a.i. L^-1^) and *S. avenae* (0.62 mg a.i. L^-1^), are comparable with the EC_50_ values reported in the susceptible populations used in similar studies, including populations confirmed to contain the homozygous susceptible *kdr*-SS allele: 0.59 mg a.i. L^-1^ in deltamethrin-susceptible *R. padi* populations (Wang *et al*., 2020, 2021). Therefore, we are confident that our detection of decreased pyrethroid sensitivity in *R. padi* population RP-5 and our range of susceptibilities and tolerance detected in our *S. avenae* populations are comparable with susceptible clones.

## Acknowledgements

This project was supported by the Alexander von Humboldt Foundation through a Postdoctoral Research Fellowship to DJL. The authors would like to thank Alexander Manentzos, Anna-Lena Heitkamp, Kristina Dauven, Eric Mühlnikel, Christoph Harms, and Antonia Pahl for assisting with aphid sampling. The authors would like to thank S Donner and M Beekman (Wageningen University & Research, The Netherlands) for kindly providing positive endosymbiont DNA samples.

## Data accessibility

The data that support the findings of this study are available from the corresponding author upon reasonable request.

## Notes

### Competing Interest Statement

The authors have declared no competing interest.

