## Supplementary material for "Common facultative endosymbionts do not influence sensitivity of cereal aphids to pyrethroids": Table S1

**Table S1: Primers used in the endosymbiont characterisation**

| **Endosymbiont target** | **Target gene** | **Primer name** | **Primer sequence** | **Concentration in mastermix** | **Estimated size (bp)** | **Multiplex experiment** |
| --- | --- | --- | --- | --- | --- | --- |
| *B. aphidicola* | 16S rRNA | 16SA1 | AGA GTT TGA TCM TGG CTC AG | 0.2 µM | 270 | Extraction test |
|  |  | Buch270R | TGC CTT GGT AGG CTA TTA CTC | 0.2 µM |  |  |
| Spiroplasma | 16S rRNA | 16SA1 | AGA GTT TGA TCM TGG CTC AG | 0.2 µM | 1500 | 1 |
|  |  | Spi500R | ATC ATC AAC CCT GCC TTT GG | 0.2 µM |  |  |
| *R. insecticola* | 16S rRNA | 16SA1 | AGA GTT TGA TCM TGG CTC AG | 0.1 µM | 840 |  |
|  |  | PAUS16SR | TCG GAC GCC ATA ACA CTA GG | 0.2 µM |  |  |
| *H. defensa* | 16S rRNA | 16SA1 | AGA GTT TGA TCM TGG CTC AG | 0.2 µM | 480 |  |
|  |  | PABS480R | GGT ATT CGC ATT TAT CGC TTC | 0.2 µM |  |  |
| *Rickettsiella spp.* | 16S rRNA | P136F | GGG CCT TGC GCT CTA GGT | 0.2 µM | 300 |  |
|  |  | P136Ric-470R | TGG GTA CCG TCA CAG TAA TCG A | 0.2 µM |  |  |
| *F. symbiotica* | 16S rRNA | PAXS_F | AGT TTG ATC ATG GCT CAG ATT G | 0.2 µM | 500 | 2 |
|  |  | PAXS_R | GCA ACA CTC TTT GCA TTG CT | 0.2 µM |  |  |
| *S. symbiotica* | 16S rRNA | 16SA1 | AGA GTT TGA TCM TGG CTC AG | 0.2 µM | 1140 |  |
|  |  | PASS1140R | TTT GAG TTC CCG ACT TTA TCG | 0.2 µM |  |  |
| *Rickettsia spp.* | 16S rRNA | 16SA1 | AGA GTT TGA TCM TGG CTC AG | 0.2 µM | 600 |  |
|  |  | Ric600R | TTT GAA AGC AAT TCC GAG GT | 0.2 µM |  |  |
| *Arsenophonus spp..* | 16S rRNA | 16SA1 | AGA GTT TGA TCM TGG CTC AG | 0.2 µM | 456 | 3 |
|  |  | Ars16S_R2 | CCT TAA CAC CTT CCT CAC GAC | 0.2 µM |  |  |

Primer details and assay from Beekman *et al.,* 2022
